## Supplementary Information for "Elastic analysis bridges structure and dynamics of an AAA+ molecular motor"

Victor Hugo Mello
*Gulbenkian Institute for Molecular Medicine (GIMM), Lisbon, Portugal*

Jiri Wald
*University Medical Center Hamburg-Eppendorf (UKE),*
*Institute of Microbial and Molecular Sciences, Hamburg, Germany*
*Centre for Structural Systems Biology (CSSB), Hamburg, Germany and*
*Deutsches Elektronen-Synchrotron Zentrum (DESY), Hamburg, Germany*

Thomas C Marlovits
*University Medical Center Hamburg-Eppendorf (UKE),*
*Institute of Microbial and Molecular Sciences, Hamburg, Germany*
*Centre for Structural Systems Biology (CSSB), Hamburg, Germany and*
*Deutsches Elektronen-Synchrotron Zentrum (DESY), Hamburg, Germany*

Pablo Sartori
*Gulbenkian Institute for Molecular Medicine (GIMM), Lisbon, Portugal*
(Dated: October 2, 2025)

### APPENDIX A: COMPARISON BETWEEN ELASTIC PSEUDOENERGY AND OTHER METRICS

In order to assess the relevance of the elastic pseudoenergy introduced in this work, we compared it against a diverse set of established metrics that have been used to characterise protein energetics, deformation, and flexibility. While each of these reference metrics is grounded in different frameworks and is associated with different spatial scales, they all serve as proxies for understanding the energetic and structural behaviour of proteins.

The first comparison involves **biophysics-based energy estimations**, where proteins are treated as systems transitioning between discrete functional states — such as active and inactive conformations — guided by free energy differences. These models, often parameterised from experimental data, provide a simplified yet informative representation of protein dynamics and energetics.

Second, we consider **force-field-based energy estimations**, which form the basis of molecular dynamics (MD) simulations. In this framework, atomic or coarse-grained interactions are encoded in empirical potential functions known as force fields, which control the time evolution of protein structures under thermal fluctuations. These simulations produce energetically informed trajectories that reflect the dynamic behaviour of proteins *in silico*.

We also explore metrics based on **Ramachandran angles**, which offer a geometric and energetic perspective on protein backbone conformation. Deviations from the well-characterised $(\phi, \psi)$ angle distributions – originally established by Ramachandran and colleagues [1] – are commonly interpreted as indicators of structural strain or steric hindrance.

Finally, we examined correlations with the **$\beta$-factor**, a metric routinely derived from X-ray crystallography and cryo-electron microscopy data. Although originally introduced to quantify atomic positional uncertainty, $\beta$-factors have been implicated in reflecting aspects of conformational entropy, local flexibility, and even functional motion, thus making them a valuable, yet indirect, probe of intrinsic structural variability [2–4].

By comparing the elastic pseudoenergy to these four distinct metrics, we aim to test whether our formulation correlates with biophysical and structural features already encoded in well-established energetic and flexibility measures.

#### Biophysics-based energy estimation

While it would be desirable to study RuvB within this framework to compare with our estimates of elastic pseudoenergy, several challenges arise. As we show in the main text, RuvB function involves coordination among subunits that themselves can adopt numerous conformations, making such energy inference difficult. Instead, we applied this approach to a cooperative model of haemoglobin conformational changes, a classical system for studying allostery and cooperativity at equilibrium that has been investigated for over a century [5].

Among the extensive literature on oxygen binding at haemoglobin's four heme-containing sites [6], a notable model is the Monod-Wyman-Changeux (MWC) model, which introduced the causal relationship between conformational changes and cooperativity [7]. Also, this model was the first to refer to distinct conformational states of the haemoglobin tetramer, T (tense) and R (relaxed), with different affinities for binding oxygen. The nomenclature remained established in the field after experimental verification of distinct conformational states. We collected examples for different types of models for the $O_2$ saturation curves of haemoglobin and show in Table I.

| **Model type** | **Reference** | **Energy difference (kcal/mol)** |
|---|---|---|
| Adair's model | Imai, 1982 [8] | 1.3 - 3.6* |
| Molecular code | Smith & Ackers, 1985 [9] | $5.9 \pm 0.2$ |
| Molecular code | Parrella et al., 1990 [10] | 7 |
| Molecular code | Huang et al., 1996 [11] | $6.3 \pm 0.2$ |
| Concerted (MWC) | Yonetani et al., 2002 [12] | 0.7 - 3.1* |
| Elastic pseudoenergy | This study ($\alpha_2\beta_2$) | $6.7 \pm 0.5$ |
| Elastic pseudoenergy | This study (2 $\alpha\beta$) | $5.2 \pm 0.4$ |

TABLE I. *Energy difference between haemoglobin R and T states from biophysical models and elastic pseudoenergy.* The values for estimated energy difference were derived from fitting experimental data to cooperative models in various experimental approaches. *Estimates of free-energy obtained from experiments in different buffer conditions.

Taking advantage of multiple structures of haemoglobin assemblies solved at T and R states [13], we computed the pairwise elastic pseudoenergy using the same approach we applied for the RuvB analysis (check the Supplementary methods for further details). To be precise, the dataset we used consists of crystallographic structures in the fully oxygenated state, or R (PDBs: 1LFQ, 1LFT, 1LFV, and 1LFY) and in the fully deoxygenated state, or T (PDB: 1LFL). The R structures display a single $\alpha\beta$ pair per unit cell, whereas the T structure has two full tetramers per unit cell. With these structures, we performed two types of pairwise comparisons: first, considering $\alpha\beta$ chains, resulting in four R states and four T states (Fig. S1A); and second, considering the full tetramers $\alpha_2\beta_2$, resulting in four R states (applying symmetry operations) and two T states (Fig. S1B). Using this approach, we calculated the elastic pseudoenergy difference of R x R pairs, T x T pairs, and T x R pairs. The advantage of using both approaches is that we can account (or not) for elastic deformations in the interface between two $\alpha\beta$ subunits. We also included the mean $\pm$ standard deviation for each group in Table I in units of kcal/mol to facilitate the comparison with the other observations.

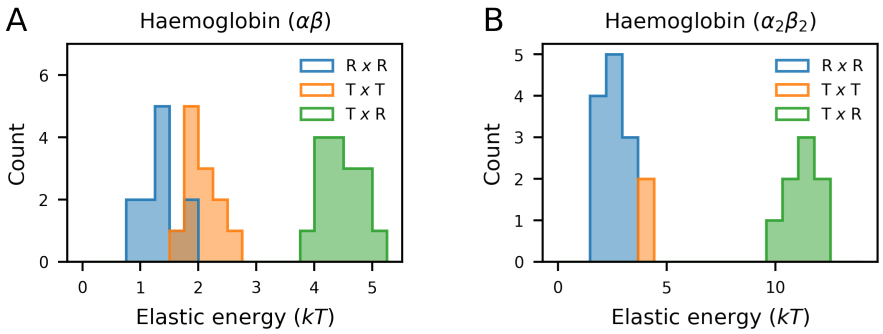


FIG. S1. *Haemoglobin elastic pseudoenergy difference from pairwise comparisons.* **A.** Comparison of individual $\alpha\beta$ pairs. With four pairs within each group, there is a total of 16 pairwise comparisons for R x R, T x T, and T x R. **B.** Comparison of $\alpha_2\beta_2$ tetramers. With four R and two T tetramers, the number of comparisons is 16 for R x R, 2 for T x T, and 8 for T x R.

Analysing the magnitude of elastic pseudoenergy in the dataset we used, we can pose interesting observations regarding haemoglobin conformational changes:

- Our results for T x R comparisons are in the same order of magnitude as those obtained with different methods and probed using very different experimental approaches. Besides, our estimate of elastic pseudoenergy falls within the equivalence range for the molecular code model. However, instead of supporting the validity of a specific model, we would rather interpret that our energy estimates match the order of magnitude of multiple biophysical models. Using strain analysis to corroborate a specific biophysical model of oxygen binding with haemoglobin requires a dedicated study for this purpose.

- Our observations reveal a consistent ordering of elastic pseudoenergy difference between types of comparison, namely, R x R < T x T < T x R. This suggests that (*a*) the energy difference between T and R states is significantly larger than those observed for T and R individually; (*b*) the R states are less variable than T states, which is in remarkable agreement with the analysis performed by the authors who solved these structures [13]. While the first observation comes with no surprise, the second is counterintuitive given the labels Relaxed and Tense. This suggests that having oxygen bound to haemoglobin has a stabilising effect on its structure.

- Comparing the same groups using only two or all four chains provides an interesting pattern: whereas R x R and T x T comparisons with $\alpha\beta$ have exactly half of the elastic pseudoenergy of $\alpha_2\beta_2$, that is not the case for T x R comparisons. This is illustrated in the last two items of Table I. This result implies that there is a significant signal of deformation localised in the interface between the two $\alpha\beta$ dimers exclusively in R $\leftrightarrows$ T transitions. This particular observation is known since 1968, when the first structures of oxyhaemoglobin and deoxyhaemoglobin were compared [14].

In summary, here we present a system where both biophysical measures of cooperative energy and structural information were available for a pair of conformational states. Using protein elasticity, we find that the elastic pseudoenergy difference between these states corresponds to the free-energy difference derived from biophysical models, within the same order of magnitude.

#### Force-field-based energy estimation

To examine how the elastic pseudoenergy calculated in this study relates to force-field-based energy estimates, we selected the all-atom force field model ff19SB [15], which is currently recommended by Amber [16], a widely used suite of programs in the molecular dynamics community. Our objective was to use this force field solely to quantify the energy of different RuvB conformations.

The protocol we followed is outlined as follows: *(i)* We converted PDB files of monomeric RuvB conformations into an Amber-compatible format using `pdb4amber`, ensuring that only residues present in all conformations were retained. *(ii)* Using the program `tleap`, we generated a simulation environment for each structure individually. We selected an implicit solvent model to prevent the initial coordinates of water molecules from influencing the energy calculations. *(iii)* We employed the program `sander` to simulate a trajectory consisting of a single iteration step. To preserve the original atomic coordinates, we deliberately avoided energy minimization, a procedure commonly used in MD protocols.

Unlike protein strain analysis, force-field calculations do not rely on a reference structure as a baseline. Consequently, we compared the elastic pseudoenergy to both the absolute value of the elastic pseudoenergy for each conformation in the cycle and the energy difference between cycle and reference conformations (see Fig. S2). We observed a significant correlation only for the energy difference, which was negative. This negative yet statistically significant correlation may be a consequence of the reference structure we chose. Because $s_0$ requires an additional ATP hydrolysis step to start the mechanochemical cycle, it might have an excess of free-energy compared to the reference structures.

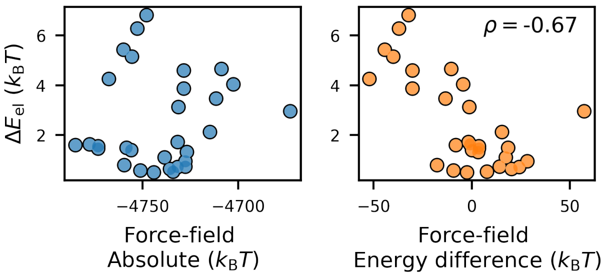


FIG. S2. *Correlation of elastic pseudoenergy and force-field derived energies.* Scatter plots show the energy values obtained for the cycle conformations. The total elastic pseudoenergy calculated from strain analysis is shown on the vertical axis, and the Amber force-field ff19SB energies on the horizontal axis. The left panel represents the absolute energy of each monomeric conformation, whereas the right panel represents the energy difference between the cycle and its corresponding reference. The strain-based elastic pseudoenergy correlates significantly with the force-field energy difference between cycle and reference ($p = 5 \cdot 10^{-5}$).

Additionally, the total energy amplitudes obtained from force-field calculations exceeded $100 k_B T$, which would correspond to an elastic model with a stiffness constant more than ten times greater than typical estimates for protein stiffness. However, this high amplitude far surpasses the non-equilibrium energy input from the ATP hydrolysis cycle and, therefore, is not consistent with the behaviour expected of a processive motor.

### Ramachandran angles

Due to the striking conservation of Ramachandran angle distributions across amino acids of the same type, the fraction of outliers is widely used as a criterion for assessing the quality of atomistic protein models [17]. Because of such relationship between conformation and energetics, we compared our inferred elastic pseudoenergy to two metrics: the Ramachandran score calculated using a standard validation tool; and the variation in Ramachandran angles between cycle and reference structures.

For the first comparison, we employed MolProbity (updated January 2024) [17], a PDB validation tool, to calculate the Ramachandran score (here denoted as $s$) for each residue. This score, derived from dihedral angles, ranks residues on a scale from 0% to 100% based on the angular distributions observed in curated high-quality structures. Lower $s$ values correspond to less populated regions of the distribution. To ensure parity with our structural comparison approach, we assigned each residue the minimum $s$-value between the cycle and reference structures.

Residues were grouped into bins ($s \leq 1\%$, $1\% < s \leq 2\%$, $2\% < s \leq 4\%$, $4\% < s \leq 10\%$, $s > 10\%$) for each amino acid type (Fig. S3). Despite limited sample sizes in lower $s$ bins, we observed a trend of increasing median elastic pseudoenergy as $s$ decreased. This suggests a detectable association between high backbone torsion and elevated elastic pseudoenergy. Glycine residues, however, deviated from this pattern, exhibiting lower elastic pseudoenergy at intermediate $s$ values. This anomaly likely stems from glycine's exceptional flexibility, due to its minimal steric hindrance (its side chain consists of a single hydrogen atom).

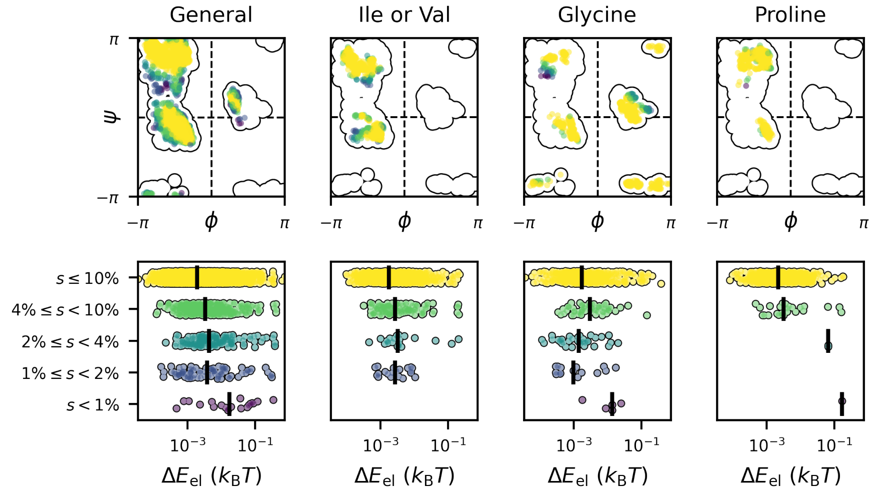


FIG. S3. *Residues near forbidden areas of Ramachandran plot tend to have higher elastic pseudoenergy.* On top, the backbone dihedral angles $\phi$ and $\psi$ are shown in the Ramachandran scatter plot colored by groups from yellow (far from forbidden regions) to purple (close to forbidden regions). Each panel represents a category of residues with specific steric constraints due to their side chain chemical composition. The contour lines draw the boundaries containing all RuvB residues in all conformations. Below, residues belonging to the corresponding category above are binned in groups according to their Ramachandran score measured with MolProbity. Black dashes mark the median of the distribution for each bin.

We next examined correlations between elastic pseudoenergy and changes in Ramachandran angles. For each residue, we computed $\Delta\phi^2 = \left(\phi_{\text{cycle}} - \phi_{\text{ref}}\right)^2$ and $\Delta\psi^2 = \left(\psi_{\text{cycle}} - \psi_{\text{ref}}\right)^2$ to quantify angular shifts.While both elastic pseudoenergy and dihedral angle differences serve as local deformation estimates, they operate at distinct spatial scales. Dihedral angles depend on the coordinates of four covalently linked backbone atoms, making them sensitive to highly localised conformational changes. In contrast, elastic pseudoenergy aggregates atomic displacements within a 9 Å radius, capturing deformations across residues that may not be adjacent in sequence.

Significant correlations emerged between elastic pseudoenergy and Ramachandran angle variations at both residue and conformation levels, particularly in their first and second moments (Fig. S4). Notably, correlations at the whole-conformation scale were far stronger than those at the residue level. This discrepancy implies that while individual residues may exhibit weak associations between deformation metrics, non-local compensatory effects in backbone

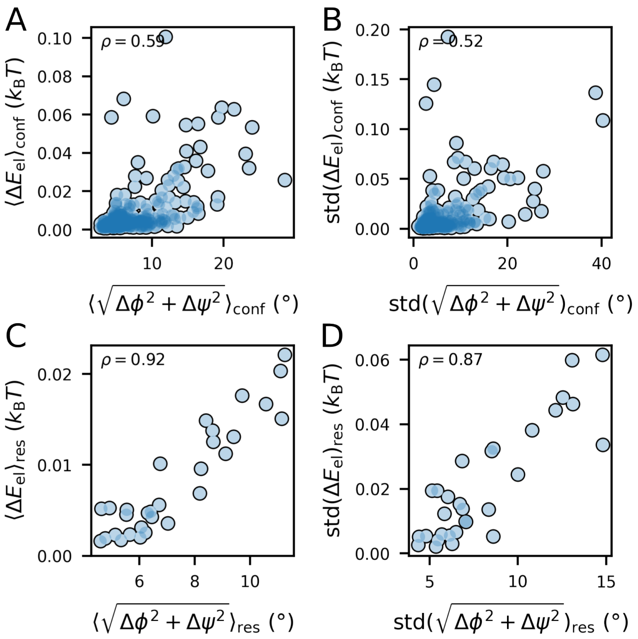


FIG. S4. *Correlations between elastic pseudoenergy and Ramachandran angle variation.* **A.** Comparison of means across conformations (correlation test, $p = 4 \cdot 10^{-30}$). **B.** Comparison of standard deviation across conformations ($p = 8 \cdot 10^{-23}$). **C.** Comparison of means across residues ($p = 4 \cdot 10^{-13}$). **D.** Comparison of standard deviation across residues ($p = 4 \cdot 10^{-10}$).

torsion collectively produce a coherent deformation profile at the global scale.

### $\beta$-factor

A comprehensive free-energy description of a protein includes both enthalpic and entropic contributions, which in principle links $\beta$-factors to the free-energy landscape of protein structures. However, caution is warranted when comparing $\beta$-factors to the elastic pseudoenergy estimates generated by our methodology. The elastic pseudoenergy calculated here does not directly account for entropic effects on atomic positions and thus primarily reflects enthalpic (internal) energy rather than the full free-energy. Moreover, there is no fundamental reason to expect a direct correlation between enthalpic and entropic forces. Additionally, regions of low structural resolution could bias elastic pseudoenergy estimates to larger values, potentially leading to spurious correlations between energy and entropy.

To explore the relationship between elastic pseudoenergy and $\beta$-factors, we computed the mean $\beta$-factor per residue (averaged across 30 conformations) and per conformation (averaged over all residues), considering the maximum $\beta$-factor between cycle and reference structures for each case. Our analysis revealed significant correlations between elastic pseudoenergy and $\beta$-factors in both the first and second moments at the residue scale, and in the first moment at the conformation scale (Fig. S5). These findings are consistent with previous studies that reported similar correlations between strain-based metrics and $\beta$-factors [18, 19]. Importantly, while positive correlations alone cannot distinguish whether elevated elastic energies arise from imprecise atomic coordinates or from regions of high conformational entropy, the collective results from this and previous analyses suggest that the elastic pseudoenergy we calculate is not simply an artifact. Rather, it robustly aligns with multiple energy-related definitions across structural scales.

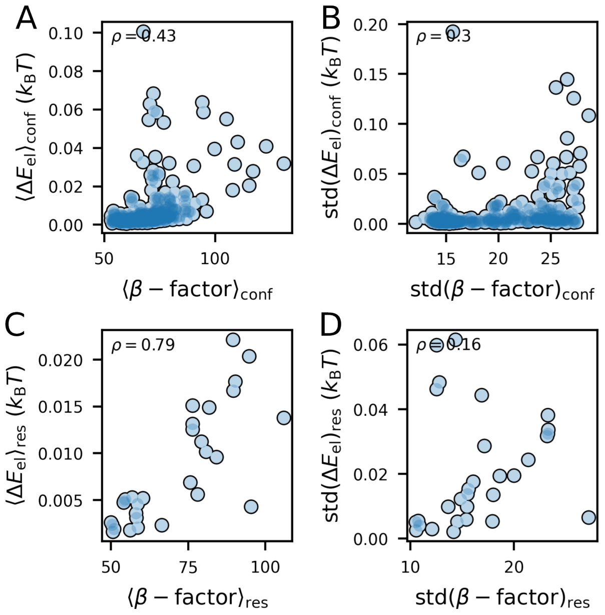


FIG. S5. *Correlations between elastic pseudoenergy and $\beta$-factor.* **A.** Comparison of means across conformations (correlation test, $p = 2 \cdot 10^{-15}$). **B.** Comparison of standard deviation across conformations ($p = 8 \cdot 10^{-8}$). **C.** Comparison of means across residues ($p = 2 \cdot 10^{-7}$). **D.** Comparison of standard deviation across residues ($p = 0.4$).

### APPENDIX B: ELASTIC MODULI OF PROTEINS

Different elastic moduli are related to various forms of deformation exerted on materials, such as shear (shear modulus), compression (bulk modulus), or one-directional extensions (Young's modulus). Most measurements of the Young modulus of proteins were obtained for polymers (summarised in [20, 21]), which might not reflect the elastic properties of single molecules due to relative displacement between protomers. These estimates range several orders of magnitude, spanning 1MPa to 10GPa. We then decided to base our stiffness constant on studies where the Young modulus was inferred using atomic force microscopy on single molecules of globular proteins. The results range from 2.5-9MPa in [22], 4-50MPa [23], 2-75MPa in [24], and 500MPa in [25]. Using a different nanorheology approach, Wang and Zocchi estimate Young's modulus of a globular protein to be around 11MPa [26].

The elastic pseudoenergy density in the Saint Venant-Kirchhoff model depends linearly on proteins' stiffness in parameters $\lambda$ and $\mu$, which are themselves functions of the Young modulus and Poisson's ratio. Because the estimates of protein stiffness vary by orders of magnitude, our estimations of elastic pseudoenergy also comprise a large uncertainty in its amplitude. The dependency of $\lambda$ and $\mu$ on Poison's ratio is non-linear and mostly flat for values close to $\frac{1}{3}$, typically assumed for proteins [25]. For our analysis, we considered $Y = 100$MPa and $\nu = \frac{1}{3}$, which led respectively to $\lambda \approx 73$MPa and $\mu \approx 38$MPa.

### APPENDIX C: LOW-DIMENSIONAL REPRESENTATION OF PROTEIN CONFORMATIONS

To perform a low-dimensional embedding of the conformational ensemble, strain is used as a measure of dissimilarity between two conformations. Specifically, we computed strain for 900 pairs of structures ($30 \times 30$), corresponding to the five hexamers in the mechanochemical cycle ($s_1, \ldots, s_5$). The strain for each pair was calculated using the PSA parameters detailed in the Supplementary Methods.

After performing all pairwise comparisons, we constructed a pairwise strain matrix $M$, where each element represents the mean strain $\langle \epsilon_3 \rangle$ across all $N$ atoms included in the analysis, $\langle \epsilon_3 \rangle = \sum_i^N \epsilon_{i,3}/N$. Since $M$ is not necessarily symmetric, we symmetrised it as $\frac{1}{2}(M + M^{\mathrm{T}})$ to obtain a suitable dissimilarity matrix for multidimensional scaling (MDS) (see Fig. S6 for a schematic overview). MDS then identifies a set of coordinates in a lower-dimensional space (here, two dimensions: $(x, y)$) for each row of the dissimilarity matrix.

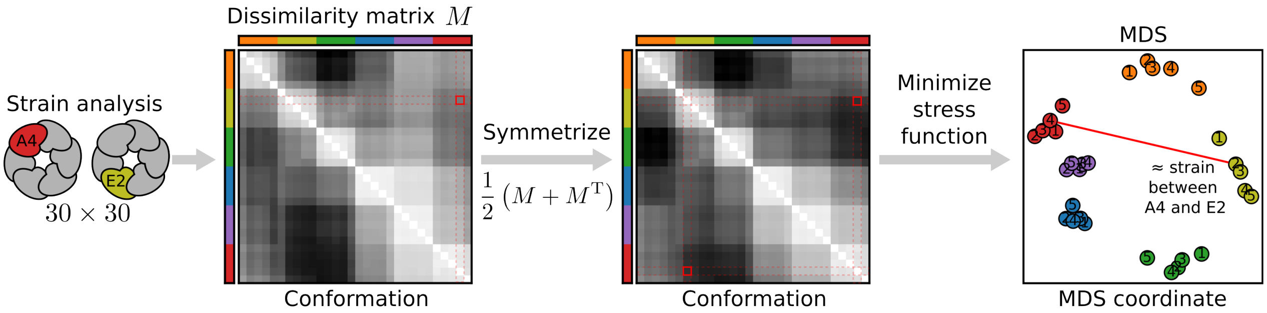


FIG. S6. *Workflow to build a low-dimensional embedding from structural comparisons.* The approach begins by calculating the local strain (or other measure of dissimilarity) for all possible pairs of conformations. In the figure, we highlight the structural comparison between conformations A4 and E2 as an example. We build a matrix $M$ with rows and columns representing particular conformations, whose elements correspond to the mean strain over the whole structure. Darker shades of gray represent high mean strain. Because such matrix is not symmetric, we apply a symmetrisation operation. Finally, we compute the MDS for the symmetrised $M$ as a dissimilarity matrix. This procedure is carried out by minimising a stress function, for which we considered an Euclidean distance on strain space.

Beyond the extensive strain $\epsilon_3$, we explored additional dissimilarity metrics to assess their ability to order the mechanochemical cycle (Fig. S7). These included the compressive strain $\epsilon_1$, the total elastic pseudoenergy $\Delta E_{\mathrm{el}}$, and local rotation angles, all derived from PSA. The local rotation angles quantify the extent to which the local atomic neighborhood undergoes rigid body rotation, as illustrated by the comparison between the rotated ellipse and the sphere in Fig. 1B. The intermediate eigenvalue $\epsilon_2$ was not considered a suitable dissimilarity metric, as it includes both positive and negative strain values (reflecting expansions and contractions). For comparison, we also constructed low-dimensional representations using root-mean-square deviation (RMSD) with three different structural alignment strategies.

The alignment approaches used in these analyses are as follows: In Fig. S7D, the entire RuvB assemblies were aligned; in Fig. S7E, individual RuvB subunits were aligned; and in Fig. S7C and F, the assemblies were aligned with an additional rigid body rotation in multiples of 60° about the central pore axis to approximately superimpose the conformations being compared.

Our results show that most metrics can capture the cyclic nature of RuvB conformations, but a clear and identifiable ordering is most apparent with $\epsilon_3$ (Fig. 2C), $\epsilon_1$ (Fig. S7A), and RMSD (Fig. S7F). Notably, both strain metrics are able to sort the cycle regardless of alignment strategy, whereas RMSD only provides a clear ordering under a specific and nontrivial alignment approach.

For completeness, we also examined how MDS places the conformations of the reference structure $s_0$ relative to the cycle conformations (see Fig. S13).

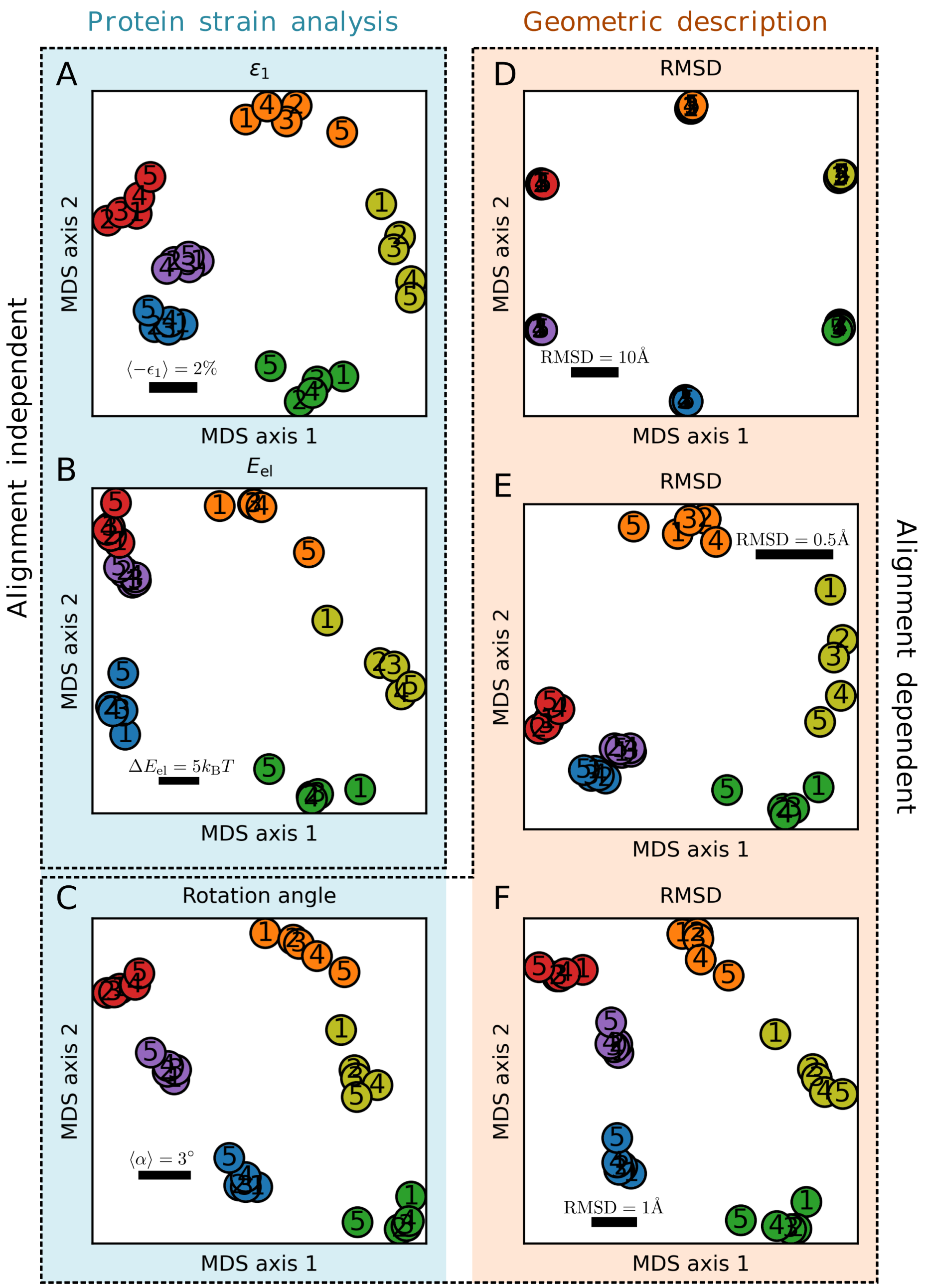


FIG. S7. *Multidimentsional scaling analyses performed with different metrics.* MDS analyses using **A.** The mean compressive strain (-$\epsilon_1$) over the protein structure. **B.** The total elastic pseudoenergy $\Delta E_{el}$ of each comparison. **C.** The mean rotation angle $\alpha$. The following panels display the root mean square deviation, RMSD, computed for three different structural alignment choices: **D.** aligning RuvB hexamers; **E.** aligning RuvB protomers; and **F** aligning RuvB hexamers and then performing rigid body rotations along the pore axis to superimpose conformations being compared.

### APPENDIX D: ELASTIC PSEUDOENERGY ANALYSES WITH ALTERNATIVE REFERENCE STRUCTURES

The quantification of RuvB's elastic pseudoenergy landscape in this study relies on the choice of a reference structure to establish an energy baseline. This choice, we stress, must be biologically justified. For RuvB, we selected a reference structure ($s_0$) solved from the same cryo-EM dataset as the cycle conformations. Critically, $s_0$ exhibits two distinguishing features: (1) it binds ATP at the terminal position adjacent to DNA, and (2) it lacks interaction with RuvA, a key cofactor essential for efficient RuvB activity. These characteristics suggest $s_0$ represents a latent, pre-engaged state of the mechanochemical cycle [27]. This interpretation is further supported by the low-dimensional projection of conformations (Fig. S13), which positions $s_0$ between $s_5$ and $s_1$.

To investigate how the choice of reference structure influences the results, we explored four additional RuvB hexameric structures as alternative references:

- 7PBQ: another potential initial state from the same EM dataset as $s_0$ but engaged with RuvA.
- 8EFV: A RuvB-DNA structure with 96% sequence coverage and 56% sequence identity to our study's constructs [28].
- AlphaFold 3 (apo): The top-ranked RuvB model (same sequence as our study) without DNA.
- AlphaFold 3 (DNA-bound): The top-ranked RuvB-DNA complex model (same sequence) [29].

The first two structures follow the same subunit labelling convention as the cycle conformations (A at the staircase top, interacting with DNA, followed by B, C, and D). In contrast, the AlphaFold models required reçlabeling to ensure consistency: the apo structure adopted a near-symmetric assembly, while the DNA-bound structure initially used divergent labels. After standardisation, we performed multidimensional scaling using extensive strain ($\epsilon_3$) as the dissimilarity metric (Fig. S8).

The results from the MDS analysis reveal distinct spatial relationships between the alternative reference structures and the cycle conformations. For example, the structure 7PBQ, much like $s_0$, clusters immediately before $s_1$ in the mechanochemical cycle, suggesting it could also serve as an initial state. In contrast, 8EFV is positioned further from the cycle conformations and does not cluster with any particular position, however, it tends to be closer to the cluster corresponding to each equivalent position than to others. The AlphaFold model generated in the absence of DNA forms a symmetric assembly, with all subunits adopting the same conformation; interestingly, this conformation resembles those found in position F (which is detached from DNA in the RuvB cycle). The DNA-bound AlphaFold model displays a more complex pattern, with some subunits matching their cognate positions (such as A and F), while the remaining subunits either form a separate cluster or do not cluster at all. Taken together, these observations indicate that only 7PBQ emerges as a plausible candidate for the initial state, closely mirroring the behaviour of $s_0$.

When constructing elastic pseudoenergy profiles using these alternative references, clear differences emerge in both the baseline and the shape of the resulting curves (Fig. S9). For all references, there is a pronounced decrease in elastic pseudoenergy upon transition to position C, except in the case of the symmetric assembly. The overall height of the energy baseline reflects how closely related the reference and cycle conformations are. However, the behaviour of the curves for positions F, E, and D varies depending on the chosen reference. For instance, the 8EFV reference exhibits two large energy barriers between F1 and F5 and between D1 and D5, while the 7PBQ reference shows a gradual increase in elastic pseudoenergy from F1 to C4. These differences highlight how the choice of reference structure can influence both the quantitative and qualitative interpretation of the elastic pseudoenergy landscape.

To further assess whether the same regions of RuvB contribute to the total elastic pseudoenergy across different conformations, we identified regions of interest as described in the main text. Consistent with previous findings, the choice of reference structure influences which residues are highlighted, resulting in varying degrees of overlap with those identified using $s_0$ as the reference. Importantly, all four regions defined in the main text were represented when using alternative references, although the specific residues included within each region differed. Regions 3 and 4 were consistently identified across all reference structures, albeit with differences in the number and identity of residues. In contrast, Regions 1 and 2 were detected in all cases except when using 8EFV as the reference, which proved to be the most dissimilar among the structures examined.

The analysis using 7PBQ as the reference, which could have been a plausible alternative for the main text, revealed two notable distinctions. First, there were fewer variable residues identified in Region 3 compared to the original analysis. Second, a novel group of residues, not previously assigned to any region, was detected. This additional group, situated within the large ATPase domain, comprises residues that interact in *trans* with Region 4 throughout the cycle. These unique features likely contribute to the observed differences in the shape of the elastic pseudoenergy landscape when 7PBQ is used as the reference. Specifically, the cycle conformations exhibit pronounced deformation in Region 3, particularly at position E, while RuvA–RuvB interactions involving Region 4 are prominent at position

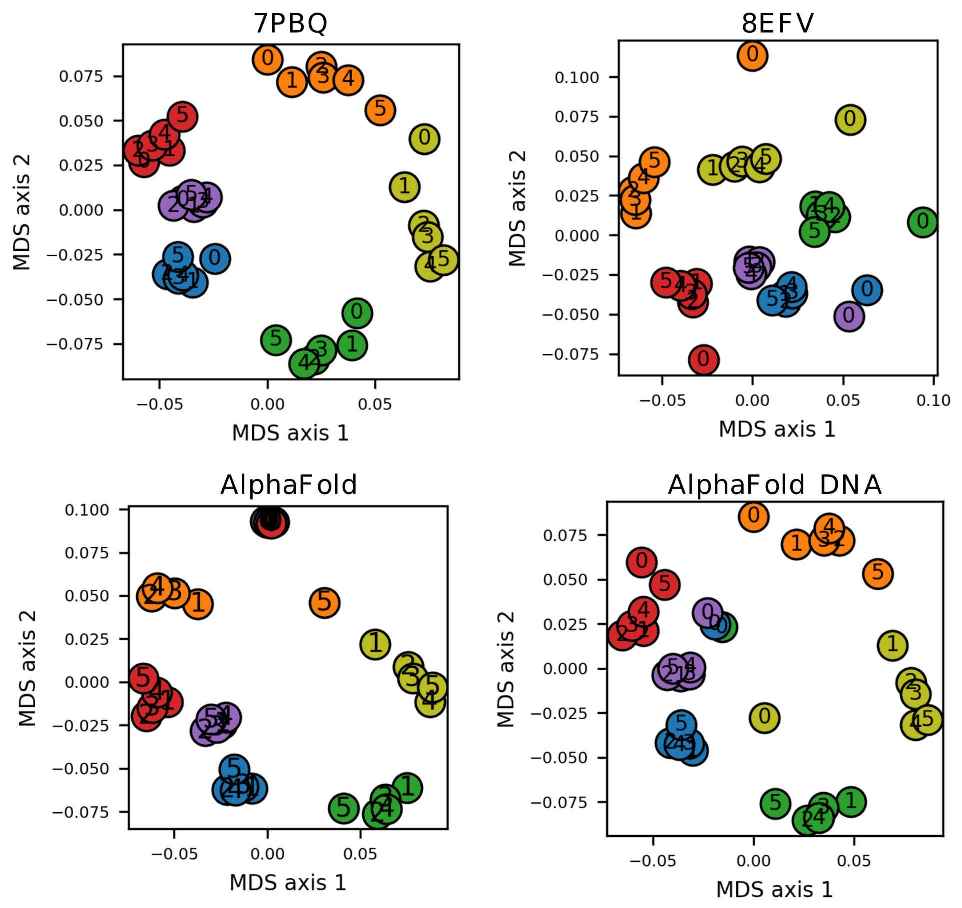


FIG. S8. *Multidimensional scaling analysis of pairwise strain with cycle and reference structures.* MDS plots using as dissimilarity metric $\langle \epsilon_3 \rangle$ between conformations. The reference conformations of each plot (shown in the title) are labelled as 0.

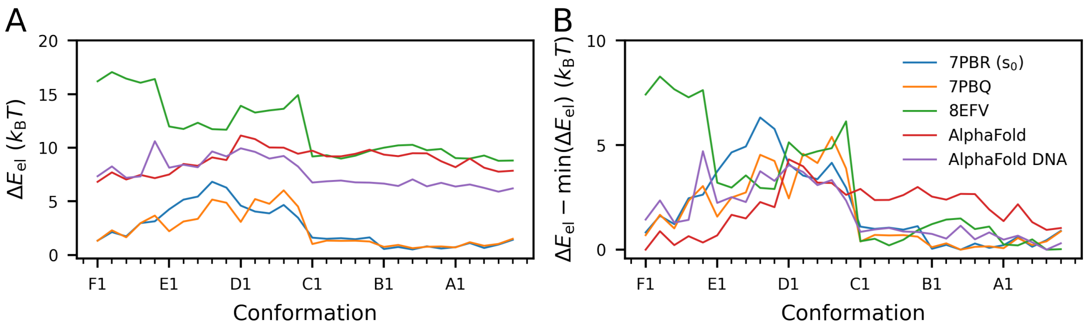


FIG. S9. *Pseudoenergy landscape of RuvB cycle using different references.* **A.** Total elastic pseudoenergy computed comparing cycle conformations to their cognate reference conformation. **B.** Amplitude of elastic pseudoenergy profile.

D. These structural distinctions align precisely with the regions of the pseudoenergy landscape where the profiles derived from 7PBQ and $s_0$ diverge.

Taken together, these results demonstrate that the choice of reference structure exerts a significant influence on the energetic characterisation of the RuvB conformational cycle. Nevertheless, the usage of our MDS-based approach provides a systematic means of evaluating and selecting biologically meaningful reference structures for further analyses.

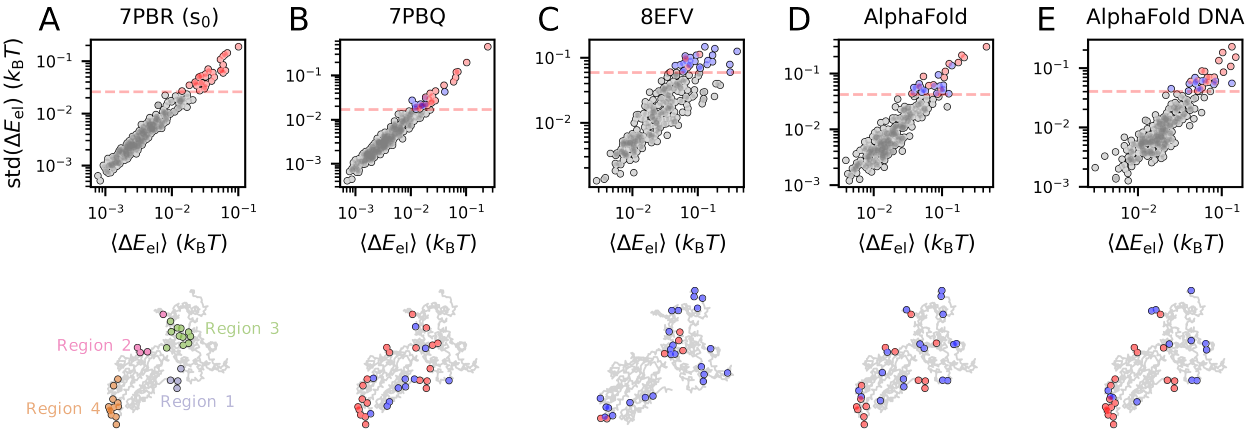


FIG. S10. *Identification of residues with most variable elastic pseudoenergy profiles using different references.* On top, we show the relationship between means and standard deviations of elastic pseudoenergy per residue. Below, we show the 10% most variable residues in a projection of a RuvB conformation. **A.** Mechanically active regions identified in the main text, using $s_0$ as reference. Residues are colored according with the scheme displayed in the main text. **B.** Most variable residues using 7PBQ as a reference shown in blue and red are compared with the mechanically active residues identified in the main text (first panel). Residues marked in red are common to the set identified for the reference $s_0$ and those in blue are not. In the following, the same procedure is repeated for references **C.** 8EFV, **D.** RuvB AlphaFold model, **E.** RuvB and DNA AlphaFold model.

### APPENDIX E: EXPLORATION ENERGY-RELATED PARAMETERS IN THE KINETIC MODEL

In this study, we develop a kinetic model that connects the microscopic conformational cycle of RuvB to its macroscopic biophysical behaviour. The model integrates three energy contributions governing conformational transitions: an energy potential, defined by a state-dependent elastic pseudoenergy landscape derived from protein mechanics; intersubunit coupling, modelled as a Kuramoto-inspired synchronisation model between subunits; and a non-equilibrium driving force, representing the chemical potential of the ATP hydrolysis cycle. In this appendix, we study how the interplay between these three energetic parameters shapes the emergent translocation speed.

We begin by highlighting the key parameters that control the amplitude of each energetic term. To explore the role of potential energy amplitude, we define a generalised potential:

$$V = V_\alpha \frac{\Delta E_{\mathrm{el}}}{\max(\Delta E_{\mathrm{el}})} \quad ,$$

where $V_\alpha$ scales the amplitude of the potential energy landscape $\Delta E_{\mathrm{el}}$ while maintaining its shape. The parameterisation used in the main text corresponds to $V_\alpha = \max(\Delta E_{\mathrm{el}}) \approx 7k_{\mathrm{B}}T$. The remaining parameters are defined as in the main text: $K$, representing the intersubunit coupling strength, and $\Delta\mu$, denoting the magnitude of the non-equilibrium drive. Figure S11 illustrates how these energetic parameters affect translocation speed.

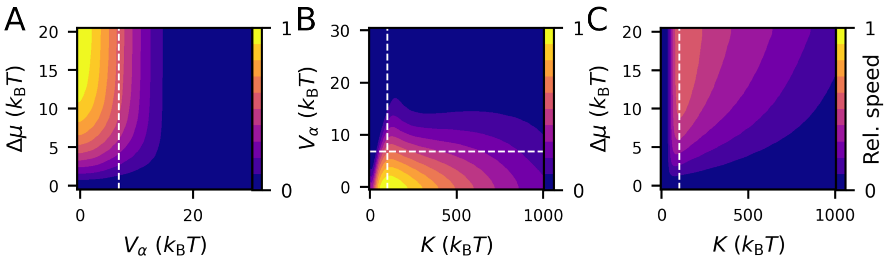


FIG. S11. *Dependence of relative processive speed on model parameters.* All heatmaps display the processive speed in colors normalised to the maximum speed achieved. **A.** Relative speed in function of potential energy amplitude and driving magnitude, at fixed coupling strength, adopted in the main text analyses ($K = 100k_{\mathrm{B}}T$). **B.** Relative speed in function of coupling strength and the potential energy amplitude, at fixed driving energy ($\Delta\mu = 20k_{\mathrm{B}}T$). **C.** Relative speed in function of coupling strength and driving magnitude, at fixed potential energy amplitude, matching main text results ($V_\alpha = \max(\Delta E_{\mathrm{el}})$). White dashed lines represent values used in the main text. All results show the mean speed of ten simulations, which were then smoothed with a Gaussian filter.

As shown in Fig. S11A, the emergent speed of the system increases monotonically with $\Delta\mu$ and decreases with $V_\alpha$. The driving energy biases the system towards an unidirectional progression, thereby increasing the motor's speed. For $K = 100k_{\mathrm{B}}T$, the speed saturates for $\Delta\mu > 15k_{\mathrm{B}}T$, as indicated by the vertical parallel level curves. Conversely, lower potential amplitudes reduce the energy barriers that individual particles must overcome to complete a full cycle, resulting in higher translocation speeds.

Figures S11B and C demonstrate that the translocation speed varies non-monotonically with the coupling strength, exhibiting a maximum around $100k_{\mathrm{B}}T$. At low coupling strengths, the trajectories of neighbouring subunits become uncoupled, preventing energy transduction between them. In the absence of coupling, particles rely solely on thermal fluctuations to cross the potential energy barrier. Conversely, at very high coupling strengths, translocation speed also declines. This decrease likely arises from a combination of finite-size effect (limited $N$) and the constraints imposed by Master-equation dynamics. As the coupling strength $K$ increases, the system must overcome a larger collective energy barrier on $E_{\mathrm{c}}$ to perform a coordinated transition from state $i$ to $i+1$ across all subunits. Such an increased barrier inflates the time scale of successful forward transitions, thereby impairing overall processive speed.

### APPENDIX F: CALIBRATION OF TIME SCALES WITH EXPERIMENTAL DATA

The kinetic model we introduced in the main text has the principle of detailed-balance as a key thermodynamic constraint, see Fig. S22 for its formulation. However, the actual form of the kinetic rates cannot be fully described by the system's energetics, as detailed-balance only dictates the behaviour of back-and-forth kinetic rates. We thus propose that a time scale variable controls the temporal dependency on kinetic rates, Eq. 8.

To calibrate such time scale, we use previously published data from two independent datasets that relate ATP concentration to translocation speeds [30]. In brief, Han and colleagues measured the [ATP]-speed curves with a bulk biochemical approach, hereby denoted as the biochemical data, and with a single-molecule experiment, resulting in what we denote biophysical data. We digitised both datasets using the web application `apps.automeris.io/wpd/`, reproduced in Fig. S12A.

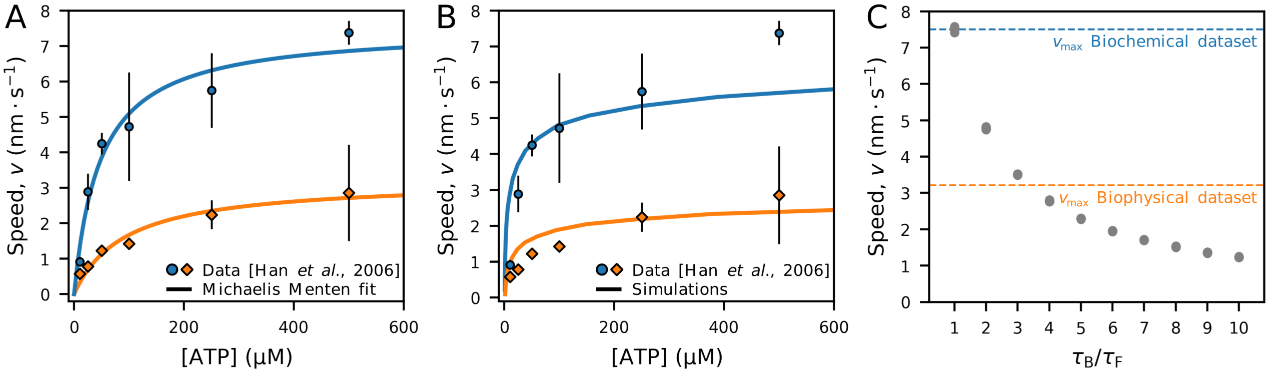


FIG. S12. *Fitting of characteristic time scale of conformational changes, $\tau$.* **A.** Reproduction of data obtained experimentally in [30], where branch migration speed $v$ is measured in two experimental approaches, namely *biochemical* and *biophysical*. **B.** Simulations adjusted to the experimental data. We sampled conformational trajectories with varying $\Delta\mu$ ranging from 0.2 to $9k_{\rm B}T$, each simulated with $n = 10$ different random seeds. **C.** Speed calculation for the biophysical dataset, with fixed $\tau_{\rm F}$ and varying $\tau_{\rm B}$ ($n = 10$ simulation replicates).

We define the time scales $\tau_{ab}$ in two forms to account for the specificities of each experimental approach. To model the biochemical dataset, we consider that all transitions have the same time scale, $\tau_{ab} = \tau_{\rm F}$. For the biophysical dataset, we model $\tau_{ab}$ as a step function to account for the slowed transitions due to the friction of the bead present in the experimental assay:

$$\tau_{ab} = \begin{cases} \tau_{\rm F} & \text{if } 0 < \theta_m, \theta'_m < \frac{2\pi}{3} \\ \tau_{\rm B} & \text{if } \frac{2\pi}{3} < \theta_m, \theta'_m < 2\pi \\ \frac{1}{2}(\tau_{\rm F} + \tau_{\rm B}) & \text{otherwise,} \end{cases}$$

with $m$ representing the molecule which underwent a state transition in $\mathbf{a} \to \mathbf{b}$. $\theta_m$ and $\theta'_m$ are the coordinates of the molecule $m$ before and after the transition, respectively. This particular choice of intervals in the reaction coordinate (0 and $\frac{2\pi}{3}$) for $\tau_{\rm F}$ and $\tau_{\rm B}$ corresponds to the RuvB conformations between free and bound. Therefore, this choice assumes that the viscous drag present in the experiment affects the transitions among DNA bound conformations. The transitions that display $\tau_{ab} = \frac{1}{2}(\tau_{\rm F} + \tau_{\rm B})$ are required to respect the symmetry condition $\tau_{ab} = \tau_{ba}$ for arbitrary neighbouring assembly states **a** and **b**.

We first calibrate $\tau_{\rm F}$, because it can be obtained directly from the biochemical dataset and our simulations. To do so, we extract the speed at saturated ATP concentration $v_{\rm max}$ fitting the biochemical dataset to the Michaelis-Menten curve $v = v_{\rm max}(1 + \frac{K_{\rm M}}{[{\rm ATP}]})^{-1}$, with $v$ the translocation speed and $K_{\rm M}$ the Michaelis constant. We then adjusted $\tau_{\rm F}$ to match the emergent translocation speed, simulating trajectories at saturated driving force ($\Delta\mu = 500k_{\rm B}T$, $n = 10$ simulation replicates), assuming that each cycle completion results in a translocation step of two DNA base-pairs, $d = 0.68$nm [27]. Next, we simulate multiple trajectories also at saturated chemical drive, with fixed $\tau_{\rm F}$ and varying $\tau_{\rm B}$, in order to find $\tau_{\rm B}$ that best reproduces $v_{\rm max}$ from the biophysical dataset, see Fig. S12C. With this approach, we obtain the time scales estimates $\tau_{\rm F} = 1.13$ms and $\tau_{\rm B} \approx 3\tau_{\rm F} = 3.39$ms.

Finally, to adjust the speed-[ATP] curves, we derive from the definition of chemical potential $\Delta\mu = \log\big([{\rm ATP}]/C\big)$, where $C$ depends on ATP, ADP, and $\rm P_i$ chemical potentials and concentrations. Then, we fit curves of processive speed simulated from a range of chemical potentials ($\Delta\mu \in [0.2, 9]k_{\rm B}T$) to the experimental data, optimising $C$ and

$K_{\mathrm{M}}$, see Fig. S12B. Our fittings reveal a poorer match to the experimental data when compared to a Michaelis-Menten fitting, as the shape of the [ATP]-speed curve for the simulations presents a sharper transition to a saturated regime. Still, the curve we obtained recovers the trend obtained from the data for an interval of $\Delta\mu$ values within the expected range of driving chemical potential observed in bacteria. For instance, the free-energy difference from the ATP hydrolysis cycle has been estimated around $\Delta\mu_{\mathrm{Phos}} \approx 47\mathrm{kJ/mol} \approx 19 k_{\mathrm{B}}T$ for *E. coli* at exponential growth [31]. Such a value should represent an upper bound to the free-energy that can be used by a molecular motor relying on ATP hydrolysis for function. This is in line with the estimates we obtain, reaching saturation on translocating speed (numerically) at $15 k_{\mathrm{B}}T$.

### SUPPLEMENTARY METHODS

#### PSA parameter choice for cycle reconstruction

We considered the following PSA parameter choice for all 900 structure pairs:

1. *Set of atoms:* heavy atoms of the main chain were selected, namely $C_\alpha$, C, N, and O.

2. *Neighbourhood radius:* for a given conformation $\alpha$, the set of atoms belonging to the local neighbourhood of $i$ was defined as $\mathcal{N}_{\alpha,i} = \{j| \left\| \mathbf{\Delta x}_i^j \right\| < r\}$, with the cutoff radius $r = 9$Å;

3. *Neighbourhood method:* to enhance the local difference between conformations, we used a linear weight method among all conformations $w_j = n(j)/N$, where $n$ is a function that counts the number of occurrences of $j \in \mathcal{N}_{\alpha,i} \forall \alpha \in \{\mathrm{F1}, \mathrm{F2}, \ldots, \mathrm{A5}\}$, in the $N = 30$ conformations. These weights represent the relative contribution of each atom in the neighbourhood to calculate the local deformation gradient.

#### PSA parameter choice for elastic pseudoenergy calculation

We considered the following PSA parameter choice to perform structural comparisons between cycle structures ($\mathrm{s}_1$, $\mathrm{s}_2$, $\mathrm{s}_3$, $\mathrm{s}_4$, $\mathrm{s}_5$) and reference ($s_0$) as follows:

1. *Set of atoms:* heavy atoms of the main chain were selected, namely $C_\alpha$, C, N, and O; the elastic pseudoenergy per residue consists of the sum of elastic pseudoenergy of its atoms.

2. *Neighbourhood radius:* for a given assembly $\alpha$, the set of atoms belonging to the local neighbourhood of $i$ was defined as $\mathcal{N}_{\alpha,i} = \{j| \left\| \mathbf{\Delta x}_i^j \right\| < r\}$, with the cutoff radius $r = 9$Å;

3. *Neighbourhood method:* to have a unified neighbourhood list per atom $i$, we considered the intersection of all neighbourhoods of $i$ in each assembly, $\mathcal{N}_i = \mathcal{N}_{\mathrm{s}_0,i} \cap \mathcal{N}_{\mathrm{s}_1,i} \cap \cdots \cap \mathcal{N}_{\mathrm{s}_5,i}$. Thus, all neighbouring atoms in $\mathcal{N}_i$ have weight $w_j = 1$;

4. *Elastic constants:* for the material properties of proteins, we considered the Young modulus $Y = 100$MPa. See the Appendix B for further information.

5. *Protin volume:*. To estimate the total van der Waals volume $V$ we used the software ProteinVolume [32] for each of the 36 RuvB conformations individually, Fig. S14C.

#### Multiple sequence alignment and conservation

Multiple sequence alignment was used to map the amino acid sequence from this study to a set of 18 bacterial RuvB sequences listed in [33]. We use the software MEGA7 to perform the multiple sequence alignment, with the gap opening penalty set to 5 and the remaining settings set to default [34].

#### Model parameters and considerations

All simulations were conducted with $\delta = \frac{\pi}{3}$ and $K = 100 k_{\mathrm{B}} T$. We chose a value $K$ that maximises translocation speed, see Appendix E. To compute the translocation speed, we considered that a translocation step of two base pairs ($\approx 6.8$Å) corresponded to a full cycle of a RuvB monomer. Also, the trajectories simulations were simulated with the initial condition $\boldsymbol{\theta}^\circ = \left[\theta_1(1), \theta_2(6), \theta_3(11), \theta_4(16), \theta_5(21), \theta_6(26)\right]$. We use this configuration because it represents an assembly state with conformations that correspond to $\mathrm{s}_1$.

**Definition of coupling energy and coupling force per state**

To calculate the coupling energy as a function of conformation, we first need to introduce the probability density of the assembly states $P(\boldsymbol{\theta})$, with $\boldsymbol{\theta} = [\theta_1, \dots, \theta_6]$. Because we are interested in describing the energetics of operational cycles, we consider here the probability density given that the initial condition $\boldsymbol{\theta}^\circ$ defined above. Therefore, we define $P^\circ(\boldsymbol{\theta}) = P(\boldsymbol{\theta}|\boldsymbol{\theta}(t=0) = \boldsymbol{\theta}^\circ)$. Other particular choices of initial state can lead to local minima of the energy landscape which are not processive. The rationale for this choice is that RuvB assembly starts the mechanochemical cycle in $s_1$. With this, we introduce the procedure to calculate $F$, the mean value of an arbitrary function $f$ evaluated in the reaction coordinates $\phi$

$$F(\phi) = \langle f(\phi) \rangle_{\boldsymbol{\theta}} = \sum_{\boldsymbol{\theta}} P^\circ(\boldsymbol{\theta}) \left[ \frac{1}{M} \sum_{m=1}^{M} f(\theta_m) \delta(\theta_m - \phi) \right] \quad ,$$

where $\delta$ is the Kronecker delta.

Using this definition, we then calculate the mean backward coupling $E_{\mathrm{c}}^{\mathrm{bwd}}$ as the mean elastic pseudoenergy $E_{\mathrm{c}}(\theta_m, \theta_{m+1})$ with the first argument fixed, and the mean forward coupling $E_{\mathrm{c}}^{\mathrm{fwd}}$ with the second argument fixed. Thus, we obtain

$$E_{\mathrm{c}}^{\mathrm{bwd}}(\phi) = \sum_{\boldsymbol{\theta}} P^\circ(\boldsymbol{\theta}) \left[ \frac{1}{M} \sum_{m=1}^{M} E_{\mathrm{c}}(\theta_m, \theta_{m+1}) \delta(\theta_m - \phi) \right] \quad ,$$

$$E_{\mathrm{c}}^{\mathrm{fwd}}(\phi) = \sum_{\boldsymbol{\theta}} P^\circ(\boldsymbol{\theta}) \left[ \frac{1}{M} \sum_{m=1}^{M} E_{\mathrm{c}}(\theta_{m-1}, \theta_m) \delta(\theta_m - \phi) \right] \quad .$$

The coupling forces are defined as the spatial derivative of the coupling energy. To compute the spatial derivative, we assume a linear relationship between conformational and physical spaces, $x_m = R\theta_m$, with the conformations arranged along a circumference of radius $R = 6$nm. Therefore, the coupling force can be computed as

$$F_{\mathrm{c}} = \frac{\partial}{\partial x} \left( E_{\mathrm{c}}^{\mathrm{bwd}} - E_{\mathrm{c}}^{\mathrm{fwd}} \right) = \frac{\partial \theta}{\partial x} \frac{\partial}{\partial \theta} \left( E_{\mathrm{c}}^{\mathrm{bwd}} - E_{\mathrm{c}}^{\mathrm{fwd}} \right) = R \frac{\partial}{\partial \theta} \left( E_{\mathrm{c}}^{\mathrm{bwd}} - E_{\mathrm{c}}^{\mathrm{fwd}} \right)$$

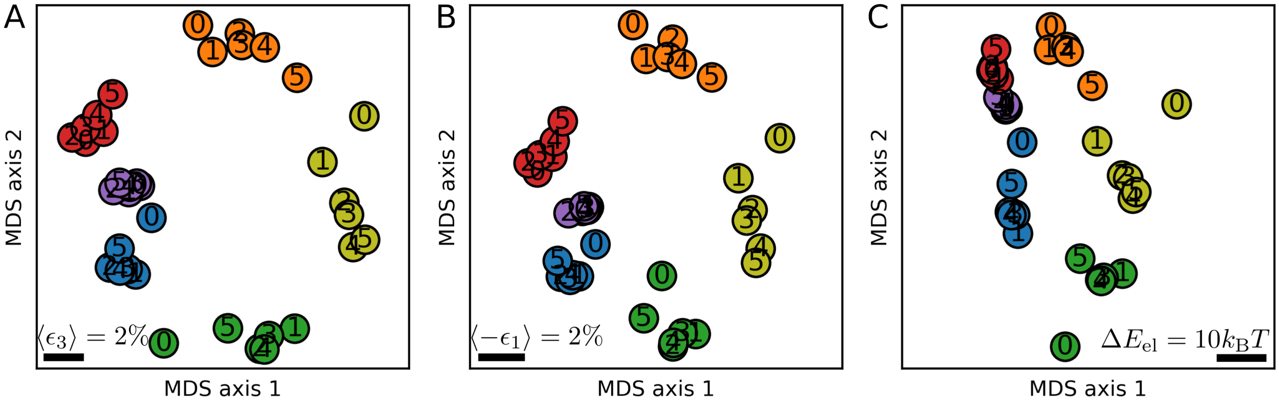


FIG. S13. *The structure* $s_0$ *in perspective of the conformational cycle.* Multidimensional scaling analysis with cycle structures and the initial state $s_0$ (mean of 100 MDS) using **A.** The mean largest eigenvalue of the strain tensor, $\langle\epsilon_3\rangle$, **B.** The mean lowest eigenvalue of the strain tensor, $\langle-\epsilon_1\rangle$, and **C.** The total elastic pseudoenergy $\Delta E_{\mathrm{el}}$ as dissimilarity measures. Despite most of the $s_0$ conformations lying before $s_1$ in clockwise order in this representation, the conformations in blue (corresponding to position C) and green (position D) are in different locations in MDS coordinates. That can be rationalised by the absence of interaction with RuvA in conformation C0 and the different nucleotide composition in position D (which harbours an ATP in conformation D0). The proximity of $s_0$ and $s_1$ for positions A, F and E suggests that $s_0$ precedes $s_1$ in order. However, the nucleotide composition of $s_0$ subunits does not accommodate it as part of the mechanochemical cycle, suggesting that a priming ATP hydrolysis in D0 is required to start the mechanochemical cycle without a position switch. Note that in the cycle conformations, hydrolysis happens only in position A, as opposed to D. Therefore, considering $s_0$ as an initial state not yet engaged in the mechanochemical cycle is a parsimonious interpretation of such MDS analyses.

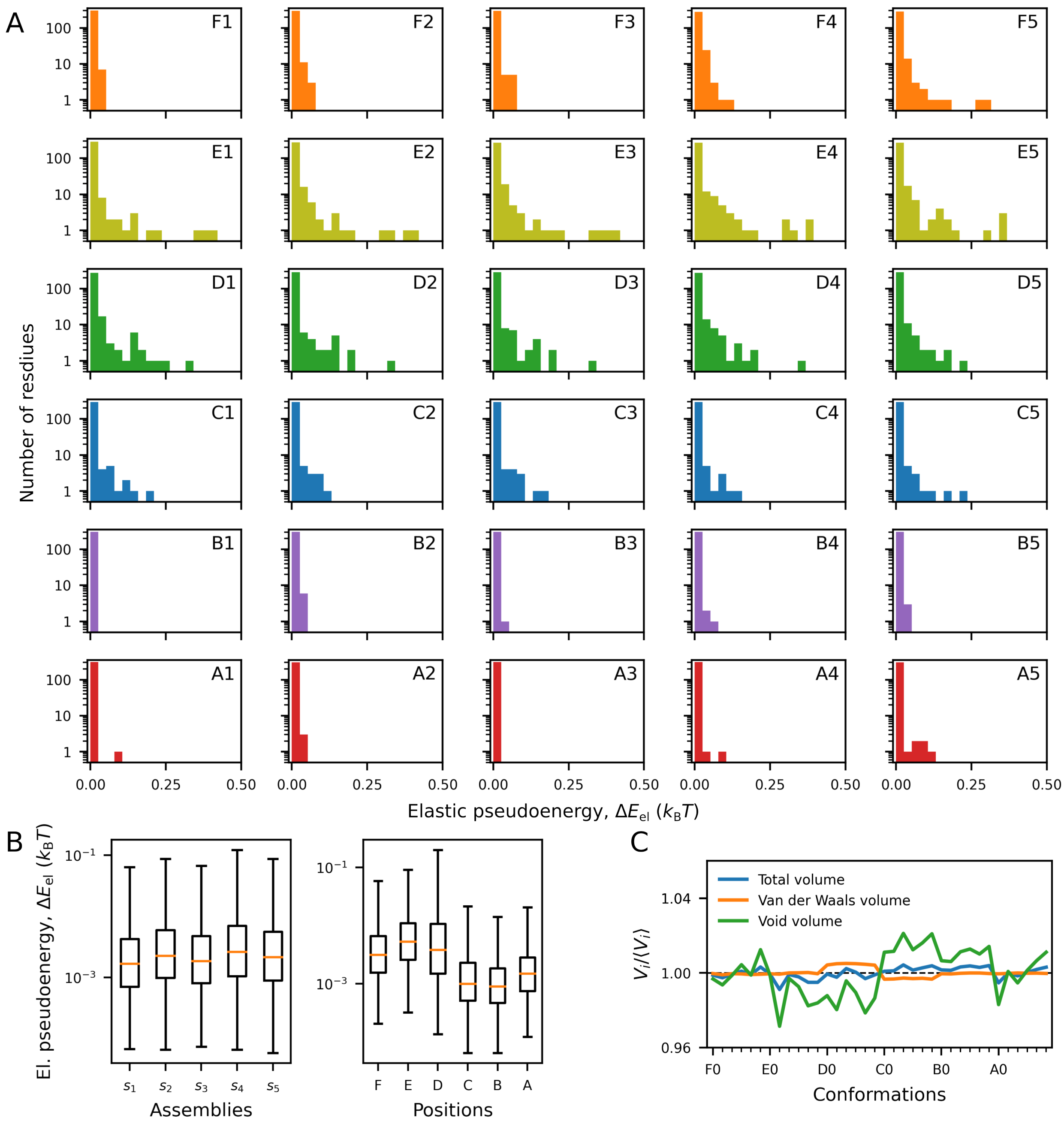


FIG. S14. *Statistics of RuvB energetics and volume.* **A.** Histogram of elastic pseudoenergies per residues in all thirty monomeric conformations. The elastic pseudoenergy distribution, depicted in log scale, shows that $\Delta E_{\mathrm{el}} < 0.01 k_{\mathrm{B}}T$ for most residues in all conformations, with heavier tails for conformations in positions E and D. **B.** Total pseudoenergy distribution among assemblies is less variable than the pseudoenergy distribution among positions. **C.** Volume per conformation displays very low variation (around 0.1%) for the van der Waals volume, which was considered for the elastic pseudoenergy calculations.

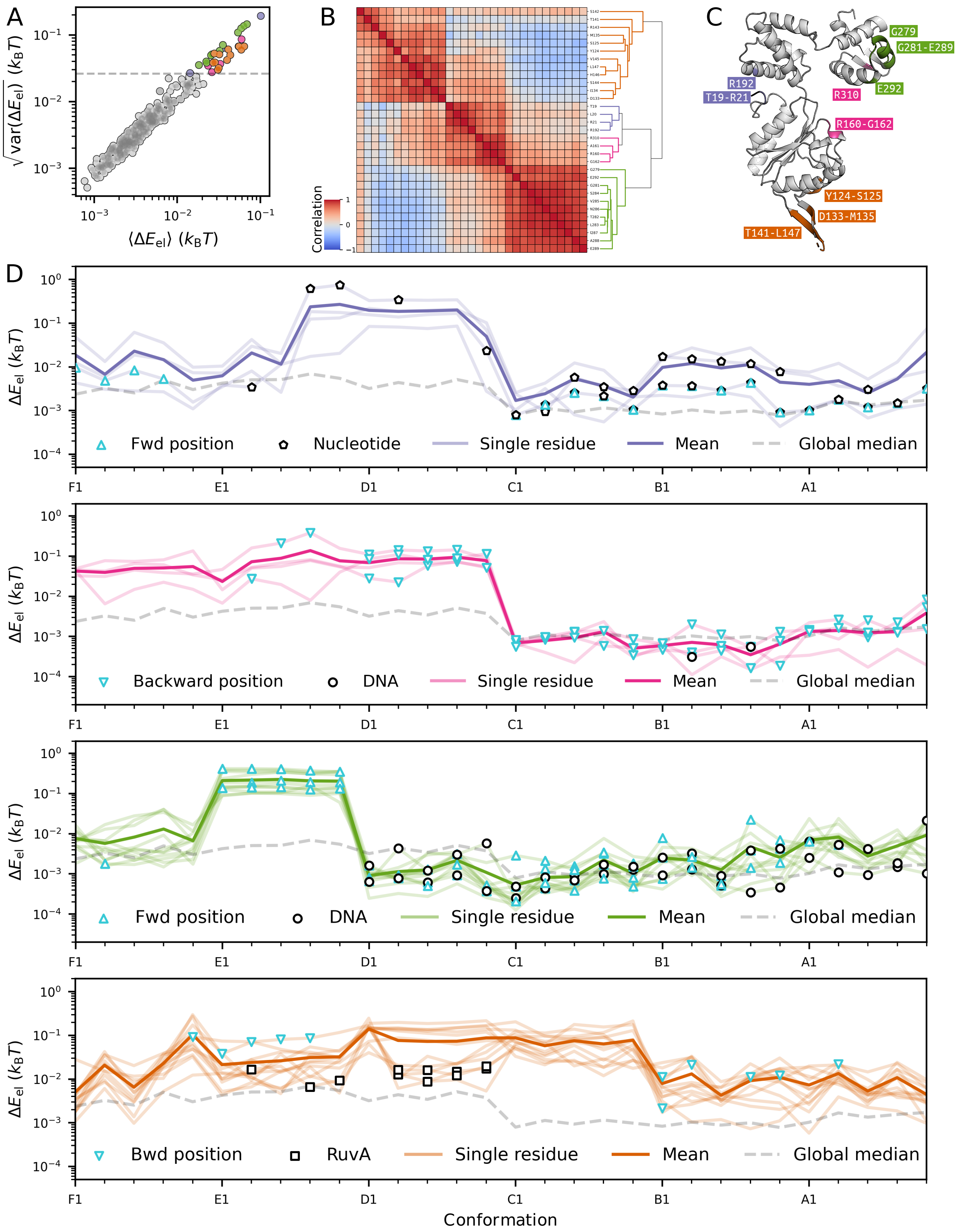


FIG. S15. *Elastic pseudoenergy profiles and intermolecular interactions performed by the mechanically active regions.* **A.** Mean and standard deviation of elastic pseudoenergy of residue trajectories are highly correlated. The dashed line defines a threshold on variance, selecting the top 10% more elastically variable residues. **B.** Correlation between elastic pseudoenergy profiles reveals clusters of residues with similar trajectories. The hierarchical clustering is based on the Euclidean distance among correlations. **C.** Mapping of spatial localisation of mechanically active regions. **D.** Elastic pseudoenergy profile of regions 1 to 4 and their specific intermolecular interactions.

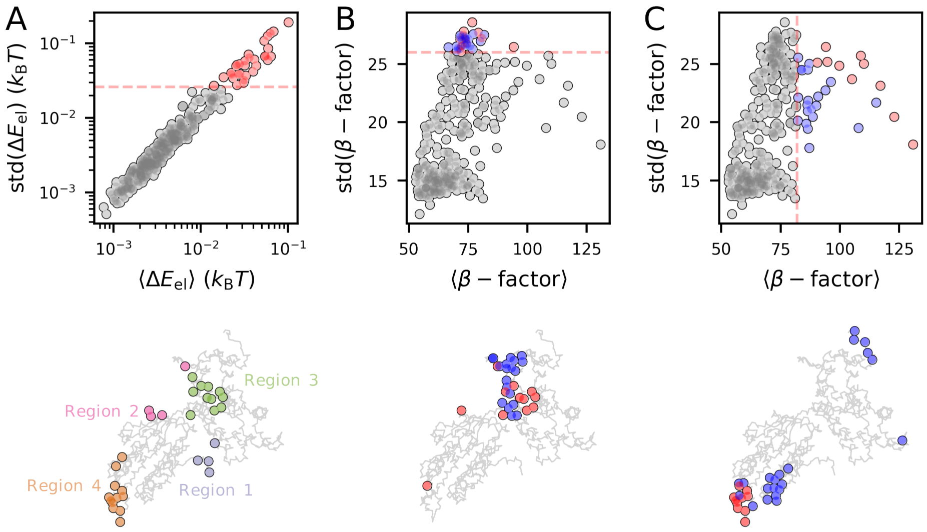


FIG. S16. *Identification of regions of interest based on elastic pseudoenergy and $\beta$-factor.* **A.** Approach used in the main text to identify regions of mechanical activity considering the top-10% residues on the standard deviation of elastic pseudoenergy. **B.** Regions of high $\beta$-factor standard deviation (top 10%). Residues marked in red agree with the elastic pseudoenergy criterion, whereas those in blue are unique to the $\beta$-factors standard deviation threshold. Residues identified above the threshold are highlighted in red and shown in a projection of RuvB monomer (below). Residues with high variability on $\beta$-factors are enriched with residues from Region 3 **C.** Regions of high $\beta$-factor means (top 10%). Residues with high mean $\beta$-factors are enriched with residues from Region 4. The results show that using $\beta$-factors does not fully recover the regions of interest identified through elastic pseudoenergy analysis, suggesting they should be interpreted as complementary rather than redundant metrics.

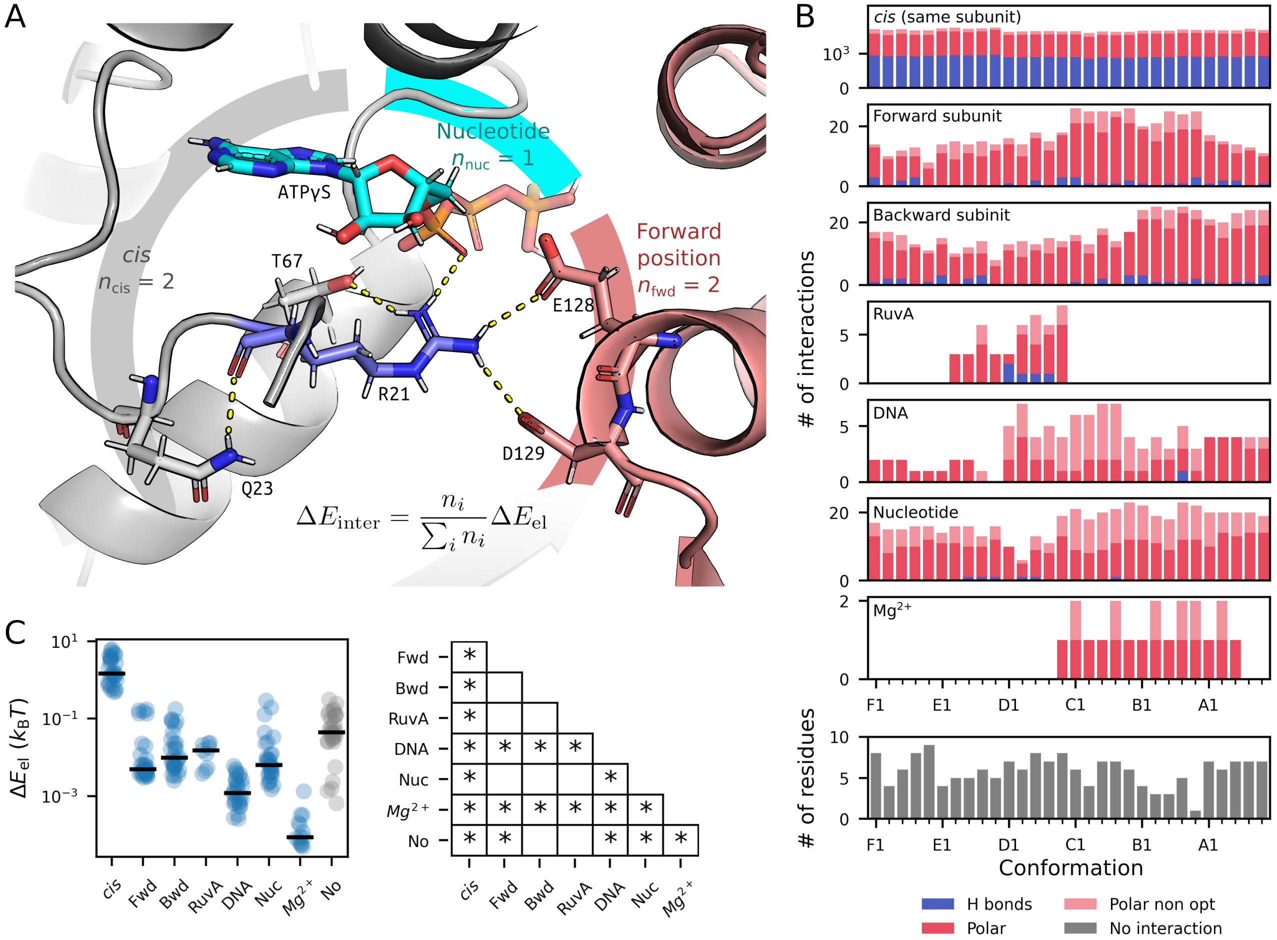


FIG. S17. *Polar interactions.* **A.** Example of polar interactions identified for R21 (in mechanically active region 1), which displays interactions in *cis*, with the forward RuvB subunit, and with the nucleotide. The elastic pseudoenergy of interaction $\Delta E_{\text{inter}}$ is calculated as the average of the elastic pseudoenergy of the residue weighted by the number $n_i$ of interactions performed. **B.** Number of polar interactions per state grouped by type of interaction. Few residues do not display any of the considered types of polar interactions. **C.** Total elastic pseudoenergy with intra and intermolecular interactions and its associated statistics. We considered only the states where there was at least one interaction. On the right, we show the statistics tested between pairs of groups (Mann-Whitney U test, * corresponds to $p < 0.05$, Bonferroni corrected).

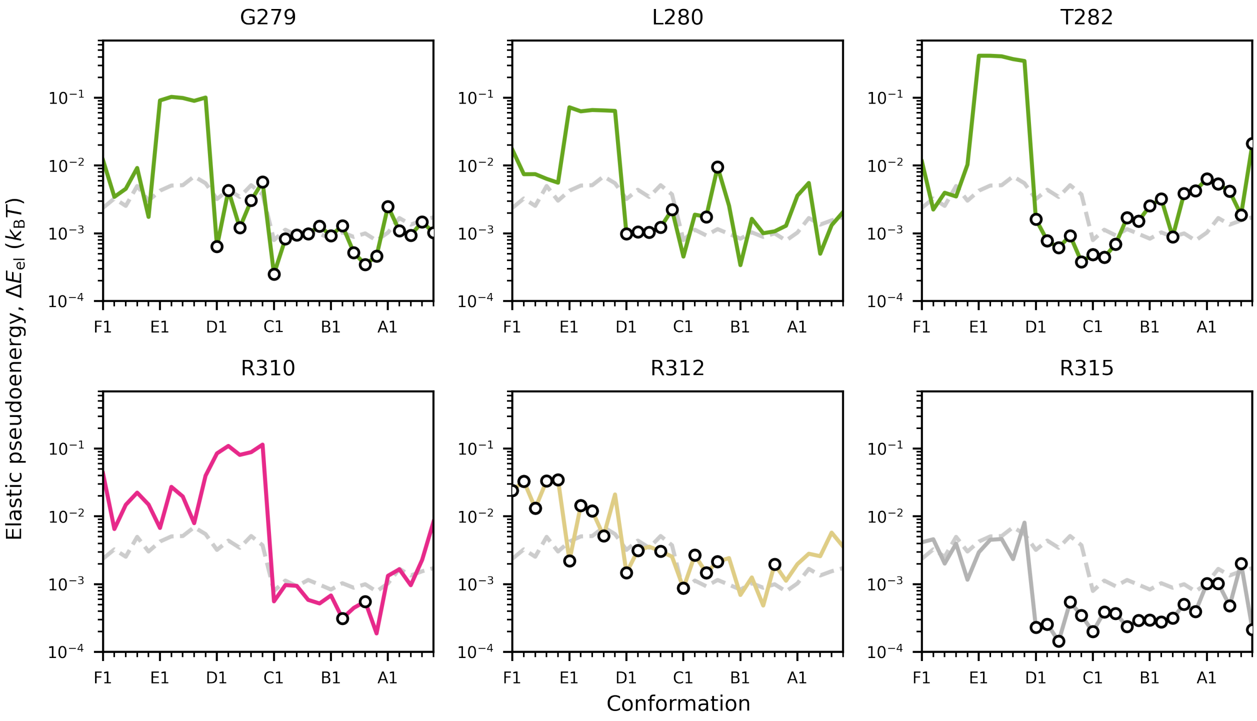


FIG. S18. *Elastic pseudoenergy and DNA interaction profiles.* Elastic pseudoenergy quantification for residues that interact with the DNA via polar interactions in at least one conformation. Circle markers indicate states with predicted interactions, and the dashed line represents RuvB's median elastic pseudoenergy profile.

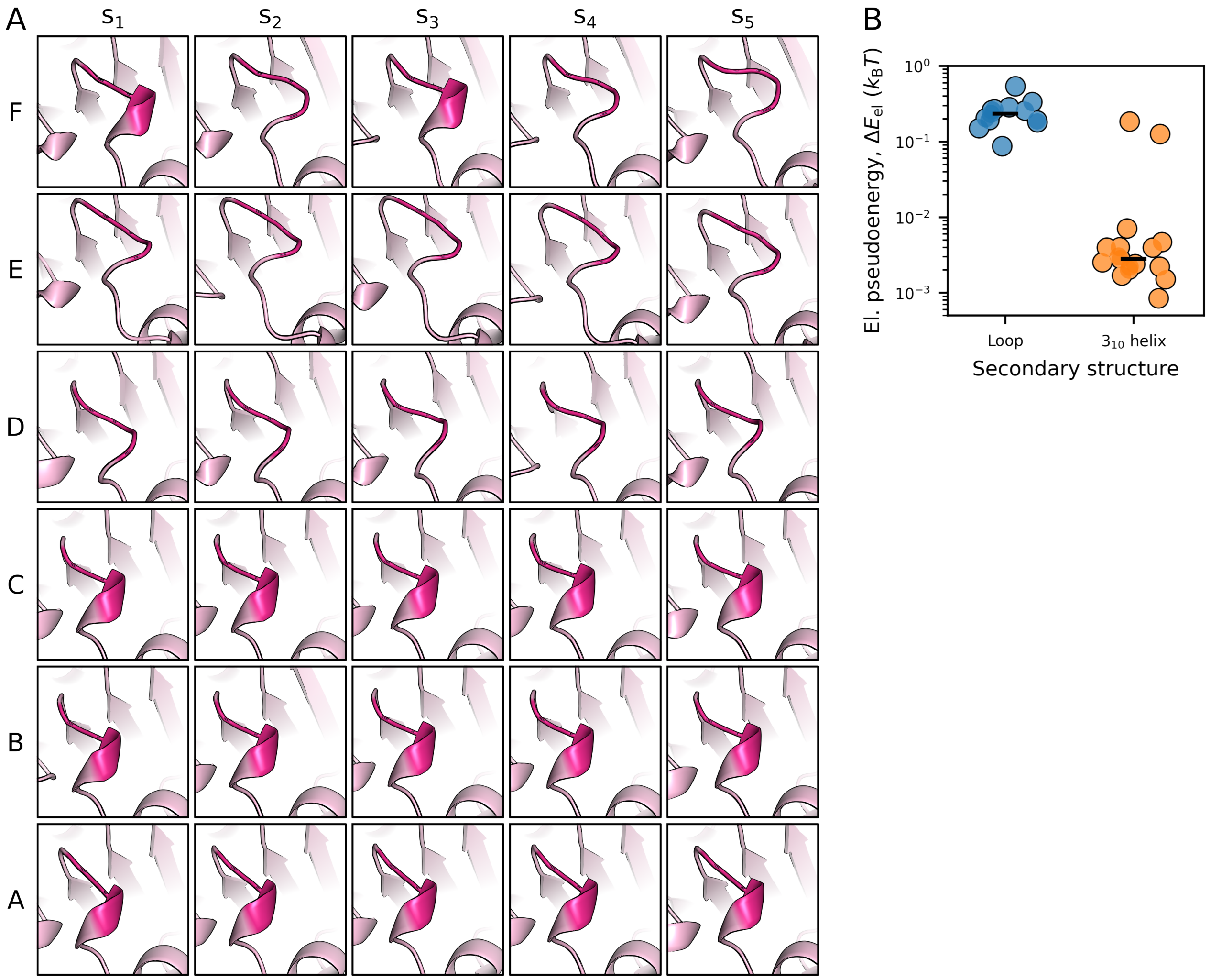


FIG. S19. *Elastic pseudoenergy content and secondary structure are correlated in Sensor 1 motif.* **A.** The residues highlighted in magenta represent part of the mechanically active region 2, found by our approach. Conformations from F to D, except for F1 and F3, were predicted to have a loop as a secondary structure, whereas conformations from C to A display a transient $3_{10}$ helix. **B.** The total pseudoenergy of residues R160, A161, and G162 is shown according with the secondary structured. The total pseudoenergy of these residues when found on loop conformation is significantly different than when on $3_{10}$ helix conformation (Mann-Whitney U test, $p < 0.05$).

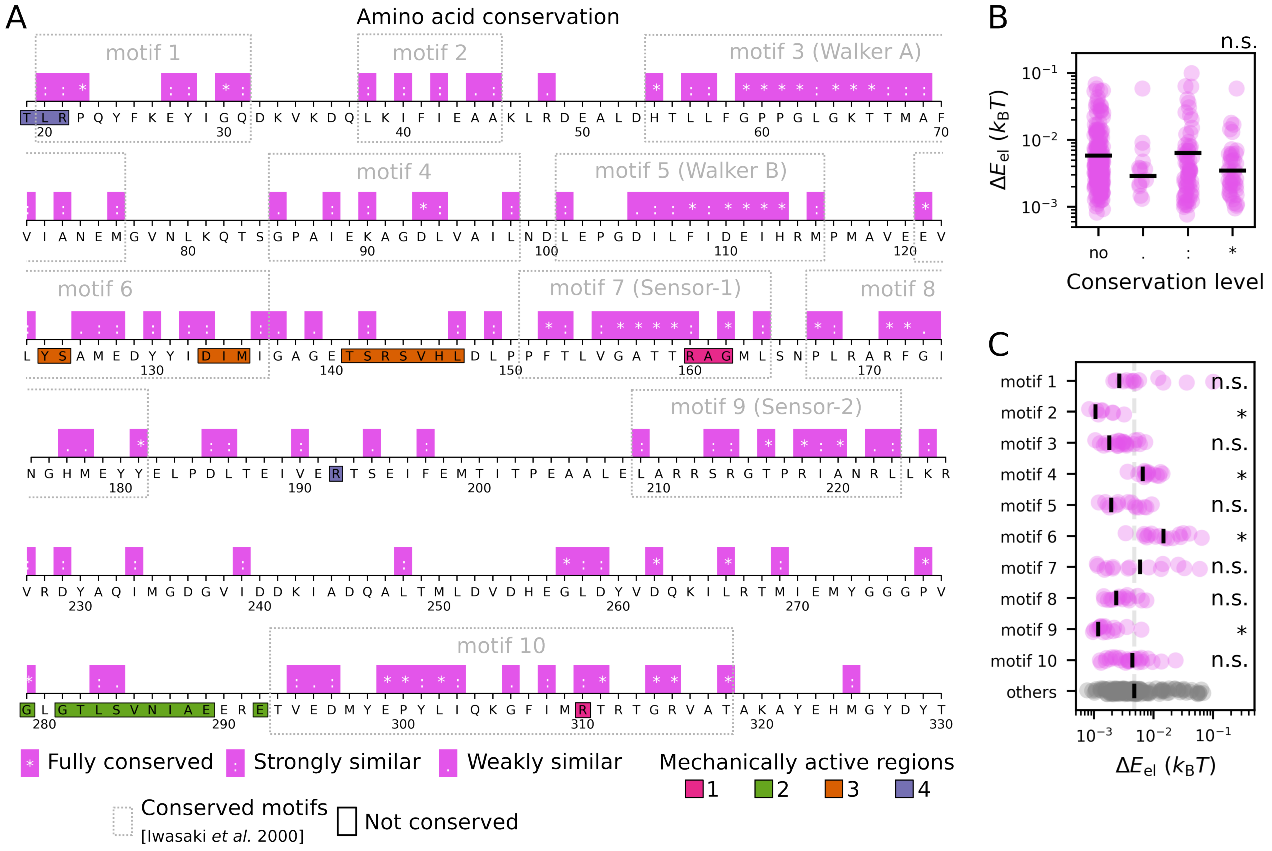


FIG. S20. *Relationship between elastic pseudoenergy and sequence conservation.* **A.** Map of amino acid conservation among RuvBs from 18 bacterial species [33] and the construct we analysed. The conserved motifs correspond to regions previously assigned by [33], to which the extent of conservation ranges from RuvB-specific to P-loop NTPases. **B.** Sequence conservation is not correlated with the mean elastic pseudoenergy per residue (Kruskal-Wallis H-test, $p > 0.05$). **C.** Mean elastic pseudoenergy of residues belonging to different conserved motifs. We found that the groups are significantly different (Kruskal-Wallis H-test, $p < 0.05$), but the pattern is not consistent across the different motifs. Motifs 2 and 9 were less energetic than non-conserved residues, whereas Motifs 4 and 6 were more energetic. Note that Walkers A and B (motifs 3 and 5) are not significantly more or less strained than non-conserved regions. (Mann-Whitney U-test for each motif *vs* other residues, Bonferroni corrected $p < 0.05$).

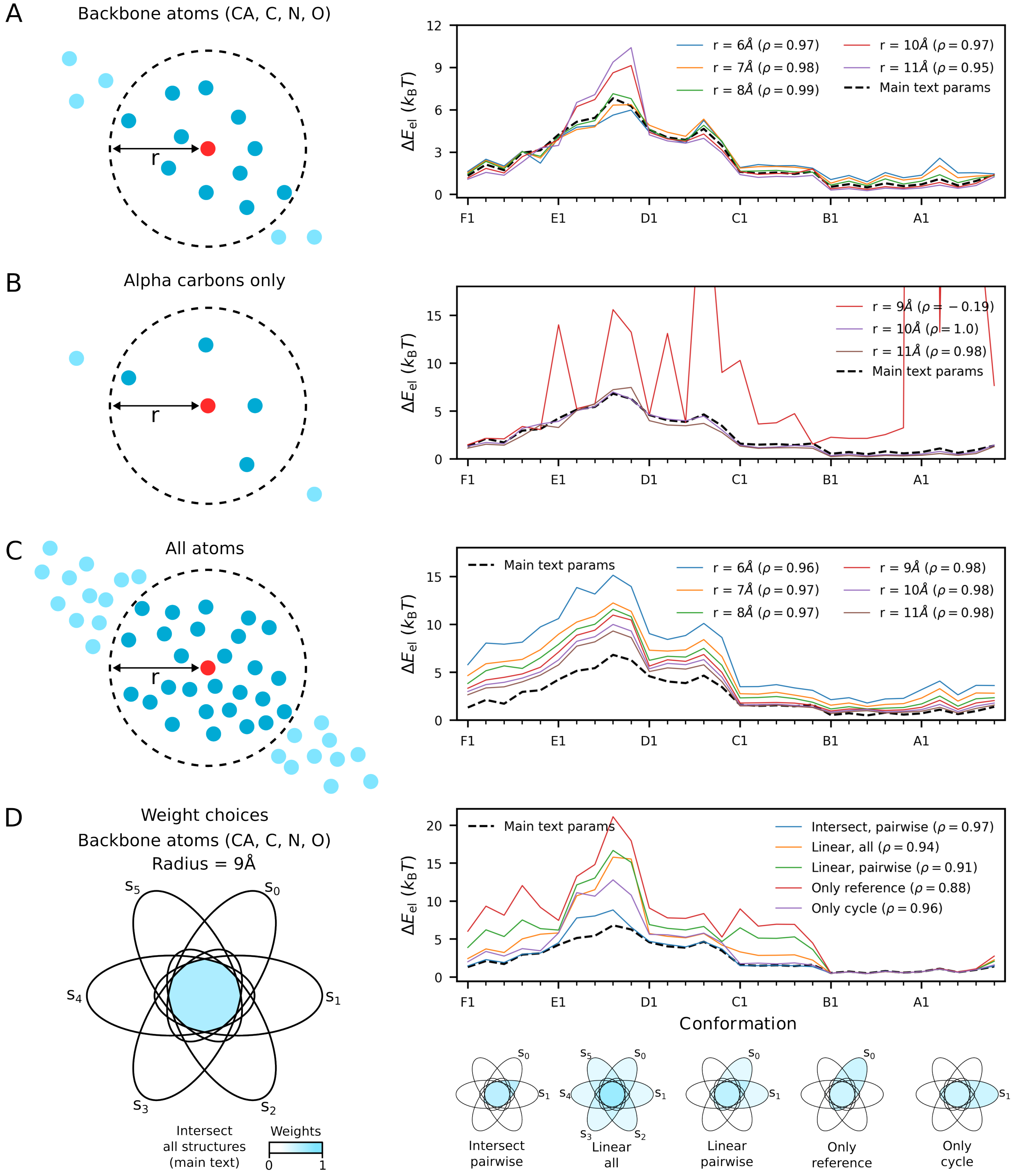


FIG. S21. *RuvB elastic pseudoenergy profile is robust to different parameter choices.* We systematically repeated the calculations for elastic pseudoenergy at different states, changing the atoms selected for the analysis, the radius of the local neighbourhood, and the method to assign weights to the atoms in the neighbourhood. We show the Pearson's correlation ($\rho$) between the pseudoenergy landscapes obtained with different parameter choices and the main text analysis (backbone atoms, $r = 9$Å, intersect all structures). Overall, the main text parameter choice agrees qualitatively and quantitatively with other choices being, in general, more conservative (typically is as deformed as or less deformed than other pseudoenergy landscapes). All profiles display a similar trend, except for alpha carbons and radius $r = 9$Å due to imprecise calculations when the number of atoms in the local neighbourhood is small. **A/B/C.** elastic pseudoenergy profiles change the radius of the neighbourhood and the atom selection criteria. For these tests, we use the same weighting method from the main text. **D.** elastic pseudoenergy profile for different weight method, using backbone atoms and radius to 9Å. The Venn diagrams represent the atoms in the neighbourhood of radius $r$ for different structures $s_i$, for the example $s_1$ vs $s_0$.

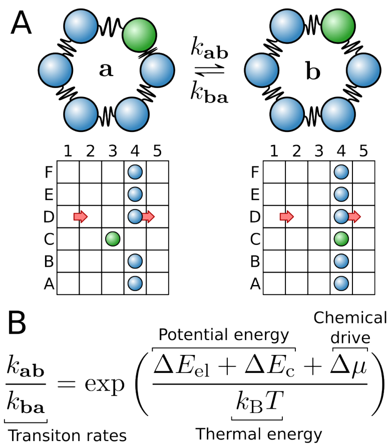


FIG. S22. *Kinetic model formulation.* **A.** Representation of a particular transition between assembly states. A RuvB assembly consists of six coupled subunits, represented by circles connected through springs. In a given hexameric state, each subunit occupies one of the 30 conformations of the mechanochemical cycle, see tables below schematics for the example states **a** and **b**. Transitions among adjacent states are controlled by the kinetic rates $k$. In the example, the green circle transitions between the adjacent conformations C3 and C4, where the other subunits remain in the same configuration. Red arrows highlight the transitions driven by the chemical potential $\Delta\mu$, which are associated with ADP release and ATP binding **B.** The detailed-balance condition enforces the ratio of back-and-forth transition rates to respect the energy difference of the assembly configurations. The driven transitions, however, are biased towards one direction.

FIG. Video 1. *Localisation and interactions of the mechanically active region 1.* The panels display the backbone of RuvB subunit and the adjacent forward subunit, with which region 1 interacts in trans. The alpha carbons of residues in region 1 are highlighted in dark purple and the residues that interact with region 1 through polar contacts are in light purple.

FIG. Video 2. *Localisation and interactions of the mechanically active region 2.* The panels display the backbone of RuvB subunit and the adjacent backward subunit, with which region 2 interacts in trans. The alpha carbons of residues in region 2 are highlighted in dark pink and the residues that interact with region 2 through polar contacts are in light pink.

FIG. Video 3. *Localisation and interactions of the mechanically active region 3.* The panels display the backbone of RuvB subunit and the adjacent forward subunit, with which region 3 interacts in trans. The alpha carbons of residues in region 3 are highlighted in dark green and the residues that interact with region 3 through polar contacts are in light green.

FIG. Video 4. *Localisation and interactions of the mechanically active region 4.* The panels display the backbone of RuvB subunit and the adjacent backward subunit, with which region 4 interacts in trans. The alpha carbons of residues in region 4 are highlighted in dark orange and the residues that interact with region 4 through polar contacts are in light orange.

[8] K. Imai, *Allosteric effects in haemoglobin.* Cambridge University Press, 1982.

[16] D. A. Case, H. M. Aktulga, K. Belfon, D. S. Cerutti, G. A. Cisneros, V. W. D. Cruzeiro, N. Forouzesh, T. J. Giese, A. W. Götz, H. Gohlke, S. Izadi, K. Kasavajhala, M. C. Kaymak, E. King, T. Kurtzman, T.-S. Lee, P. Li, J. Liu, T. Luchko, R. Luo, M. Manathunga, M. R. Machado, H. M. Nguyen, K. A. O’Hearn, A. V. Onufriev, F. Pan, S. Pantano, R. Qi, A. Rahnamoun, A. Risheh, S. Schott-Verdugo, A. Shajan, J. Swails, J. Wang, H. Wei, X. Wu, Y. Wu, S. Zhang, S. Zhao, Q. Zhu, T. E. I. I. I. Cheatham, D. R. Roe, A. Roitberg, C. Simmerling, D. M. York, M. C. Nagan, and K. M. J. Merz, “AmberTools,” *Journal of Chemical Information and Modeling*, vol. 63, pp. 6183–6191, oct 2023.

[17] V. B. Chen, W. B. Arendall III, J. J. Headd, D. A. Keedy, R. M. Immormino, G. J. Kapral, L. W. Murray, J. S. Richardson, and D. C. Richardson, “*MolProbity*: all-atom structure validation for macromolecular crystallography,” *Acta Crystallographica Section D*, vol. 66, pp. 12–21, jan 2010.

[18] M. R. Mitchell, T. Tlusty, and S. Leibler, “Strain analysis of protein structures and low dimensionality of mechanical allosteric couplings,” *Proceedings of the National Academy of Sciences*, vol. 113, no. 40, pp. E5847—-E5855, 2016.

[19] E. Weinreb, J. M. McBride, M. Siek, J. Rougemont, R. Renault, Y. Peleg, T. Unger, S. Albeck, Y. Fridmann-Sirkis, S. Lushchekina, J. L. Sussman, B. A. Grzybowski, G. Zocchi, J.-P. Eckmann, E. Moses, and T. Tlusty, “Enzymes as viscoelastic catalytic machines,” *Nature Physics*, 2025.

[20] M. Guthold, W. Liu, E. A. Sparks, L. M. Jawerth, L. Peng, M. Falvo, R. Superfine, R. R. Hantgan, and S. T. Lord, “A comparison of the mechanical and structural properties of fibrin fibers with other protein fibers,” *Cell biochemistry and biophysics*, vol. 49, pp. 165–181, 2007.

[21] J. Howard, *Mechanics of Motor Proteins and the Cytoskeleton.* Sunderland, MA: Sinauer, 1 ed., 2001.

[27] J. Wald, D. Fahrenkamp, N. Goessweiner-Mohr, W. Lugmayr, L. Ciccarelli, O. Vesper, and T. C. Marlovits, “Mechanism

of AAA+ ATPase-mediated RuvAB–Holliday junction branch migration," *Nature*, vol. 609, no. 7927, pp. 630–639, 2022.

[28] A. D. Rish, Z. Shen, Z. Chen, N. Zhang, Q. Zheng, and T.-M. Fu, "Molecular mechanisms of Holliday junction branch migration catalyzed by an asymmetric RuvB hexamer," *Nature Communications*, vol. 14, no. 1, p. 3549, 2023.

[29] J. Abramson, J. Adler, J. Dunger, R. Evans, T. Green, A. Pritzel, O. Ronneberger, L. Willmore, A. J. Ballard, J. Bambrick, S. W. Bodenstein, D. A. Evans, C.-C. Hung, M. O'Neill, D. Reiman, K. Tunyasuvunakool, Z. Wu, A. Žemgulytė, E. Arvaniti, C. Beattie, O. Bertolli, A. Bridgland, A. Cherepanov, M. Congreve, A. I. Cowen-Rivers, A. Cowie, M. Figurnov, F. B. Fuchs, H. Gladman, R. Jain, Y. A. Khan, C. M. R. Low, K. Perlin, A. Potapenko, P. Savy, S. Singh, A. Stecula, A. Thillaisundaram, C. Tong, S. Yakneen, E. D. Zhong, M. Zielinski, A. Žídek, V. Bapst, P. Kohli, M. Jaderberg, D. Hassabis, and J. M. Jumper, "Accurate structure prediction of biomolecular interactions with AlphaFold 3," *Nature*, vol. 630, no. 8016, pp. 493–500, 2024.
